# Introducing Large Language Models to Human-Based Etymological Classification in Zooplankton

**DOI:** 10.1101/2025.05.08.652882

**Authors:** Haruto Sugeno, Keito Inoshita, Kota Nojiri, Takumi Taga

## Abstract

We benchmarked LLM-based against expert epithet-etymology labelling for 989 zooplankton species, finding high concordance except in culturally nuanced names, confirming LLMs’ value as rapid screening aids.

## Introduction

Scientific names of organisms represent the fundamental units for species identification and classification. These are indispensable, particularly in taxonomy, ecology, and biodiversity research (Pat and Nicolas, 2020). These names are assigned based on standardized rules established by the International Code of Zoological Nomenclature (ICZN) and the International Code of Nomenclature for algae, fungi, and plants (ICN), enabling consistent species identification (ICZN, 1999; ICN, 2024). In recent years, with the advancement of molecular techniques such as Next-Generation Sequencing (NGS), the discovery of new species and re-evaluation of existing classifications have progressed rapidly, making the scientific name an increasingly important common information base for researchers worldwide (**Stevens** et al., 2019). However, the origins of many species names are unclear due to challenges such as stagnation in the archiving of past literature and the challenge of interpreting etymologies, as the relevant descriptions are often written in a wide range of languages, making them difficult to access and analyze systematically.(Richard, 2000). Traditionally, identifying the etymology of a scientific name involves focusing on the species epithet and carefully examining the original descriptions and related literature to infer naming intentions from the content (Macêdo et al., 2023).

This method, however, requires considerable time and effort, making it unsuitable for large-scale analysis. These issues are particularly evident in taxonomic groups with high diversity and frequent new species descriptions (Macêdo et al., 2023). Large language model (LLM), which learns from vast amounts of textual data and incorporate various forms of knowledge including taxonomic names, are expected to automate and incorporated various forms of knowledge, including taxonomic names, are expected to automate the etymological labeling of scientific names (Inoshita et al., 2025). However, studies applying LLM to etymology labeling have so far been limited to certain arthropod groups (Nojiri et al., 2025). This study, therefore, evaluates the effectiveness of automating etymological inference of species epithets in zooplankton using LLM. Additionally, the dataset used in this study has been etymologically labeled by multiple taxonomists, making it well-suited for evaluating the applicability and accuracy of LLM-based analysis (Macêdo et al., 2023). This approach enables us to capture temporal trends in naming practices and to examine how etymological categories have shifted over time. By evaluating these changes, we can also assess whether large language model (LLM) is capable of detecting shifts in cultural context reflected in species epithets. By doing so, we aim to facilitate a systematic understanding of species epithet etymology and promote the organization and practical application of taxonomic knowledge.

## Materials and methods

### Dataset

We used the open dataset “From pioneers to modern-day taxonomists: the good, the bad, and the idiosyncrasies in choosing species epithets of rotifers and microcrustaceans” published by Macêdo et al. (2023) (Macêdo et al., 2023). The dataset contains scientific nomenclature information on zooplankton species described between 1758 and 2021, and includes highly reliable records reviewed by taxonomic experts. Since it also includes names described before the establishment of the ICZN, it is useful for analyzing historical changes in naming trends. Additionally, scientific names derived from personal names are classified by whether they honor men or women. The distinction enables the investigation of gender imbalance in scientific names. To supplement the dataset, we verified original papers using SiBBr (Sistema de Informação sobre a Biodiversidade Brasileira, 2025), GBIF (Global Biodiversity Information Facility, 2025), and Google Scholar to add accurate publication years. The analysis included 989 species for which publication years could be obtained.

### Statistical Analysis

We aggregated the frequency of each category (Morphology, Ecology & Behavior, Geography, People, and Culture) from both manual and LLM classifications and visualized naming trends using bar graphs. To quantitatively assess the agreement between manual and LLM classifications, Pearson’s correlation coefficients (r) were calculated for each category. Additionally, to quantify the differences, mean absolute error (MAE) was computed for each category. Chi-square tests were conducted to evaluate the differences in category assignment between the two classification methods. Time-series trends in category frequency were modeled using generalized additive models (GAMs) to assess how naming patterns changed over time and whether trends were consistent between methods. All statistical analyses were conducted in R version 4.4.1 (R Core Team, 2024), with data import handled by the readr package (Wickham & Miller, 2023), data wrangling and transformation using dplyr (Wickham et al., 2024) and tidyr (Wickham, 2023), and GAM time-series modeling implemented via the mgcv package (Wood, 2017). Correlation coefficients and mean absolute error (MAE) were calculated using base R’s stats package.

## Results and discussion

The time required for LLM-based classification was 14 minutes, with an average of 0.85 seconds per species. A bar graph comparing category composition ratios by method revealed similar proportions for People and Geography categories between manual and LLM classifications but a notably higher proportion in the Culture category for LLM classification (Fig. 1). To evaluate the concordance between human-based and LLM-based classification, Pearson’s correlation coefficients and MAE were calculated for the five categories (Morphology, Ecology & Behavior, Geography, People, and Culture). High concordance was observed for Morphology (r = 0.978, MAE = 0.474), People (r = 0.946, MAE = 0.202), and Geography (r = 0.918, MAE = 0.156). In contrast, the Culture category showed lower concordance (r = 0.497, MAE = 0.601), indicating discrepancies between the two methods.

**Fig. 1.**
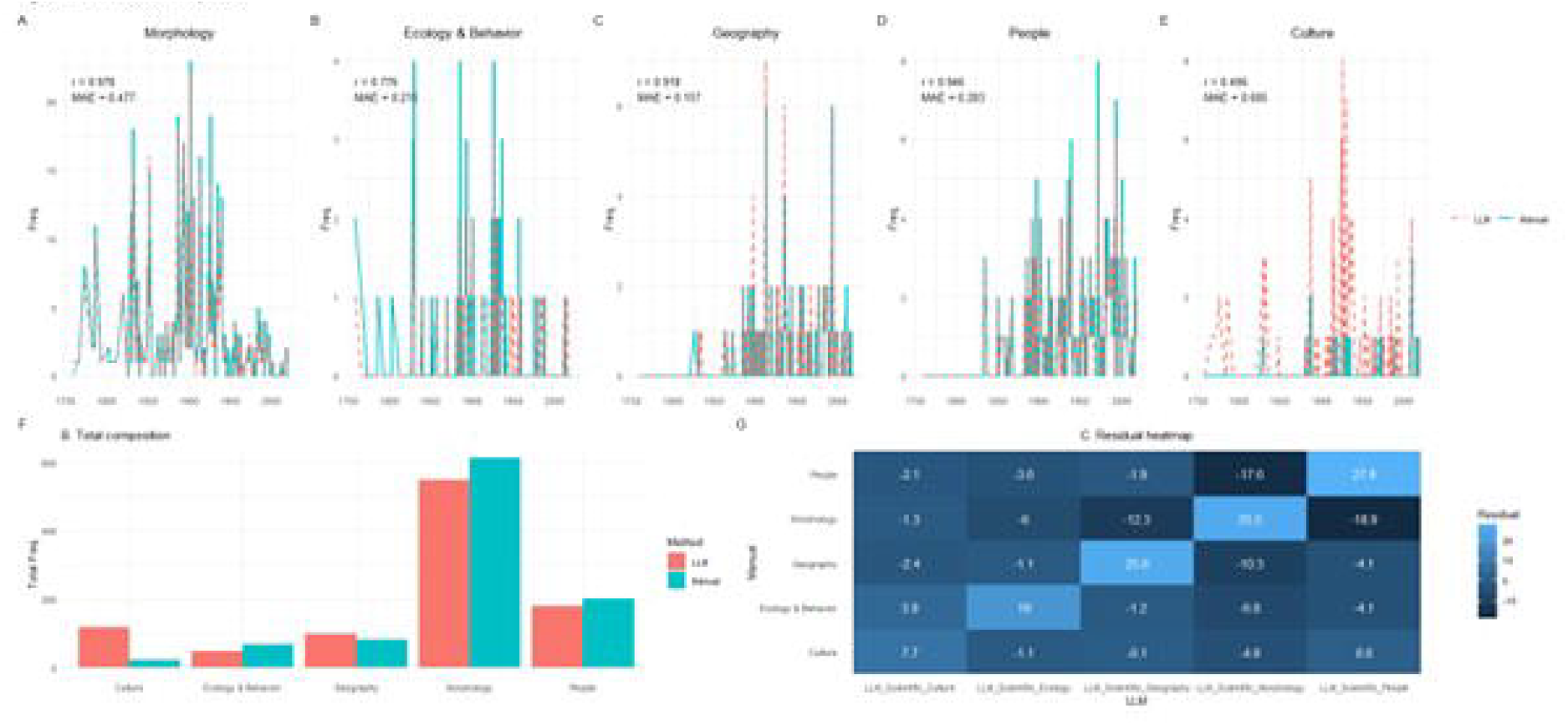
Comparison of manual and LLM epithet classifications. Yearly frequencies for five categories (A–E), total composition (F) and Pearson residual heat-map (G).

To determine whether there was a statistically significant difference between LLM and manual classifications, a chi-square test was performed, revealing a significant difference (X^2^ = 1953.7, df = 16, p < 0.001). Standardized residuals from the chi-square test showed that LLM tended to over assign the Culture category (Culture→Culture, residual = +7.70), while epithets derived from their Morphology were frequently misclassified into other categories, especially Morphology→People (residual = −18.9) and Morphology→Ecology & Behavior (residual = −6.03). Cases of overclassification of epithets derived from geographical feature into Morphology were also observed (Geography→Morphology, residual = −10.3). People→People (residual = +27.9) had the highest concordance, suggesting that LLM is highly accurate in identifying names derived from personal names. These results exceed the statistical threshold of |residual| > 2, indicating systematic bias between the two classification methods by category.

Time-series trends of category frequency in both manual and LLM classifications were compared using GAMs (Fig. 2). Both methods shared commonalities and differences in temporal trends. Morphology-based names were prevalent from the late 19th to mid-20th century but have declined in recent years.

**Fig. 2.**
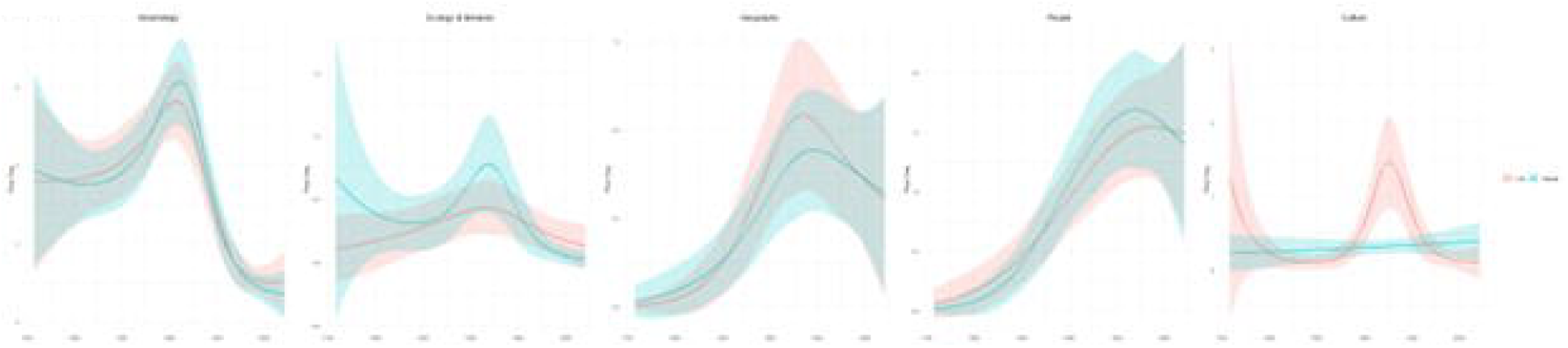
GAM-smoothed temporal trends (mean ± SE) for each epithet category, contrasting manual and LLM classifications.

The epithets based on People and Geography increased in the latter half of the 20th century, with similar trends observed for both methods. Culture-based names showed consistently low frequency overall but tended to be slightly overpredicted by LLM.

GAM smoothing showed discrepancies in Culture and Morphology, while the overall trends were consistent between LLM and manual classification.

This study compared human-based classification and LLM-based automated classification of species epithet etymologies in zooplankton and evaluated the concordance and systematic differences between the two methods.

From the Pearson’s correlation coefficients and MAE, the LLM-based classification showed high concordance with manual classification for Morphology, Geography, and People categories.

Particularly for names derived from People, the LLM demonstrated extremely high concordance (r = 0.946), reflecting the model’s strong performance in extracting proper nouns and understanding their meanings. On the other hand, the Culture category showed low correlation (r = 0.497) and a tendency toward misclassification, suggesting that interpretations of cultural or metaphorical expressions may differ between humans and LLM.

The chi-square test and standardized residual analysis revealed that the LLM had systematic biases toward certain categories. Specifically, the model tended to over assign the Culture category (+7.70 residual) while underestimating Morphology and People categories. In the Morphology category, many epithets were misclassified into other categories, suggesting that the LLM may be inclined to interpret etymologically ambiguous terms as morphological features.

Time-series analysis using GAMs showed that overall trends were similar between LLM and manual classifications. For example, Morphology-based naming was predominant in the 19th century, while naming based on Geography and People increased from the 20th century onward. The ability of LLM to replicate such long-term trends suggests that its understanding of semantics reflects temporal context to some extent, demonstrating the potential for applying natural language processing to academic research.

Nonetheless, there are some limitations to LLM-based automatic classification. First, the model showed overclassification for context-dependent categories such as culture and metaphor. Second, classification results may be influenced by biases in the training data. For instance, if certain cultural terms or naming patterns are overrepresented in the training corpus, predictions may be skewed.

Additionally, LLM does not provide confidence scores for outputs, making it difficult to manage uncertainty in ambiguous classifications (Xiong, 2024).

Despite these limitations, the findings suggest that LLM-based automatic classification can partially replace and, in many cases, effectively complement manual classification, depending on the research goals and data scale. Especially in reconstructing large taxonomic systems or analyzing geographic and temporal trends in naming practices, LLM can offer rapid and consistent screening as a supplementary tool. Future developments should consider a “human-in-the-loop” approach, where humans collaborate in the calibration and interpretation of LLM outputs (Mosqueira-Rey, 2023).

## Conclusions

This study demonstrates that large language models can effectively automate etymological classification of zooplankton species epithets, achieving high concordance with expert human labels for morphology, geography, and personal-name categories. However, LLMs tend to overpredict culturally nuanced or metaphorical categories, indicating inherent limitations in capturing semantic subtleties. As such, LLM-based tools are best deployed as rapid, large-scale screening aids, complemented by targeted human-in-the-loop validation. Future work should focus on fine-tuning models with domain-specific corpora, incorporating uncertainty quantification, and developing hybrid workflows to fully leverage automated classification in taxonomic and biodiversity research.

## Acknowledgements

Although no external collaborators were involved in this study, I am deeply grateful for the uninterrupted flow of coffee, the serenity of late nights, and the flashes of inspiration (and their disappearance) that supported the writing process.

## Data Archiving

The dataset used in this study is available from Macêdo et al. (2023) and related websites (https://sibbr.gov.br/, https://www.gbif.org/ja/).

